# Ecologically relevant thermal fluctuations enhance offspring fitness: biological and methodological implications for studies of thermal developmental plasticity

**DOI:** 10.1101/2020.06.23.167593

**Authors:** Joshua M. Hall, Daniel A. Warner

## Abstract

Natural thermal environments are notably complex and challenging to mimic in controlled studies. Consequently, our understanding of the ecological relevance and underlying mechanisms of organismal responses to thermal environments is often limited. For example, studies of thermal developmental plasticity have provided key insights into the ecological consequences of temperature variation, but most laboratory studies use treatments that do not reflect natural thermal regimes. While controlling other important factors, we compared the effects of naturally fluctuating temperatures to commonly used laboratory regimes on development of lizard embryos and offspring phenotypes and survival. We incubated eggs in 4 treatments – 3 that followed procedures commonly used in the literature, and one that precisely mimicked naturally fluctuating nest temperatures. To explore context-dependent effects, we replicated these treatments across two seasonal regimes: relatively cool temperatures from nests constructed early in the season and warm temperatures from late-season nests. We show that natural thermal fluctuations have a relatively small effect on developmental variables but enhance hatchling performance and survival at cooler temperatures. Thus, natural thermal fluctuations are important for successful development and simpler approximations (e.g. repeated sine waves, constant temperatures) may poorly reflect natural systems under some conditions. Thus, the benefits of precisely replicating real-world temperatures in controlled studies may outweigh logistical costs. Although patterns might vary according to study system and research goals, our methodological approach demonstrates the importance of incorporating natural variation into controlled studies and provides biologists interested in thermal ecology with a framework for validating the effectiveness of commonly used methods.

## INTRODUCTION

Temperature has great potential to explain variation across biological and ecological scales. Consequently, considerable effort is given to measuring thermal variation in nature and replicating this variation in controlled environments (i.e. laboratory or field manipulations; e.g. Hagstrum and Hagstrum, 1970; Folguera et al., 2011; Mitchell et al., 2015; Xing et al., 2019). Such studies uncover mechanisms that link thermal variation to important ecological and evolutionary phenomena in wild populations (Worner, 1992; Georges et al., 1994); however, the success of such work may depend heavily on the ability to precisely measure temperatures in the field and accurately replicate them in a controlled study (Faye et al., 2016; Carter et al., 2018). This is challenging since temperature varies spatially and temporally at different scales (Colinet et al., 2015; Hall and Warner, 2018; Tiatragul et al., 2019; Xing et al., 2019). Such complexity results in a myriad of context-dependent thermal effects on biological processes (Liu et al., 1995; Stilwell and Fox, 2005; De Jong et al., 2010; Bannerman and Roitberg, 2014; Manenti et al., 2014; Warner and Shine, 2011; Buckley et al., 2017) that may contribute as much to phenotypic variation as genetic factors (Noble et al., 2018). Due to technological or logistical constraints, researchers cannot perfectly replicate field temperatures, and therefore, use less complex manipulations that partially correlate with natural conditions (e.g. constant mean temperatures). Regardless, ecologists strongly advocate for recreating real-world thermal variation in the laboratory (Folguera et al., 2011; Bowden et al., 2014; Greenspan et al., 2016; Burggren, 2018; Mickley et al., 2019) because this can enhance our understanding of biological phenomena (Carter et al., 2018). However, replicating real-world temperatures has practical and statistical challenges (Colinet et al., 2015; Burggren, 2018). For example, incubators that reproduce field temperatures in a near-perfect way may be prohibitively costly or require custom-built equipment (Greenspan et al., 2016; Mickley et al., 2019). Moreover, increasing thermal complexity of treatments makes it difficult to interpret findings, replicate studies, and synthesize results across the literature (e.g. meta-analysis). Given these challenges, it is critical to understand how increasing levels of thermal complexity influence results under various contexts (e.g. across temperature).

Studies of reptiles have contributed greatly to our knowledge of the ecological and evolutionary significance of thermal variation (see Warner et al., 2018; Refsnider et al., 2019). Because most oviparous reptiles (excluding archosaurs) provide no parental care, developing embryos are often subjected to wide variation in temperature, which can substantially influence fitness-relevant phenotypes of embryos and hatchlings (Janzen, 1993; Pearson and Warner, 2018; Booth, 2018). Therefore, reptiles are often used as indicator species when assessing thermal effects of global change (Telemeco et al., 2009; Tiatragul et al., 2020; Hall and Warner, 2018). For these reasons, reptile developmental plasticity is of broad interest in ecology and evolution (reviewed by Du and Shine, 2015; Noble et al., 2018; While et al., 2018; Refsnider et al., 2019; Du et al., 2019) and has great potential to advance our understanding of thermal biology using experimental approaches (Warner et al., 2018). Early studies consisted of measuring nest temperatures with a thermometer and incubating eggs at constant mean temperatures (e.g. Licht and Moberly, 1965), which may poorly reflect natural embryo environments for many species (Booth, 2018) but were logistically simple and served as the foundation for future work. Technological advances (e.g. temperature loggers, programmable incubators), have allowed researchers to precisely record nest temperatures and utilize complex incubation treatments in the laboratory (e.g. Tiatragul et al., 2020). For example, temperature loggers can be programmed to record hourly, daily, and seasonal thermal variation inside nests (Pearson and Warner, 2018). Moreover, researchers can purchase or build incubators capable of near-perfectly replicating fluctuations in nest temperature (Greenspan et al., 2016) and associated stochastic temperature changes (e.g. heat waves) that typify real thermal environments (Hall and Warner, 2018; Pearson and Warner, 2018; Carter et al., 2018).

These technological and methodological advances have greatly increased our understanding of the mechanisms that underlie developmental plasticity and its ecological and evolutionary relevance (Georges et al., 2005; Warner and Shine, 2011; Bowden et al., 2014; Carter et al., 2018). For example, because developmental rate increases with temperature, embryos spend a greater portion of development at warmer vs cooler temperatures when nest temperatures fluctuate. In addition, natural thermal fluctuations frequently exceed the optimal developmental range of temperatures, whereas constant temperatures, even with similar means, remain within this range throughout development (Les et al. 2009). As a result, a host of phenotypes differ between eggs incubated at fluctuating vs constant temperatures, demonstrating the importance of replicating natural temperatures (Booth, 2018). However, most studies still use constant temperature treatments (While et al., 2018). Considering the potential benefits of more accurately reproducing real-world thermal environments along with the logistical and empirical costs, researchers should (but rarely do) invest great effort into determining the most appropriate level of thermal complexity to use in their research.

Our goal is to determine the effects of using various levels of thermal complexity in studies of developmental plasticity and how these effects change under different contexts (e.g. seasonal temperature differences). We used temperature data collected from lizard nests across a broad reproductive season to create 3 commonly used incubation methods and compared their effects on development with those of naturally fluctuating temperatures. We replicated each treatment across two seasonal temperature regimes: relatively cool vs relatively warm temperatures that typify nests early and late in the season, respectively. We incubated eggs under these conditions and measured a host of embryo and hatchling phenotypes. This unique 2 (seasons) by 4 (incubation methods) full-factorial design allows us to assess the context-dependent effects of incubation methods that vary substantially in thermal complexity. Our work evaluates the importance of replicating real-world temperatures in studies of thermal ecology and evolution. Because many taxa exhibit thermal developmental plasticity and inhabit thermally complex environments, our results have implications that span various evolutionary and ecological scales.

## MATERIALS AND METHODS

### Lizard collection and husbandry

The brown anole (*Anolis sagrei*) is a subtropical lizard native to islands in the Caribbean, but is naturalized in Florida, USA (Figure 1). Females lay a single-egg clutch once every 4-12 days from March to October (Hall et al., 2020). Eggs are deposited in shallow nests (1-4 cm deep) or laid on the soil surface beneath leaf litter or cover objects (e.g. rocks, logs) across a diversity of habitats; thus, embryos are subjected to a wide range of mean temperatures and thermal fluctuations during development (Pearson and Warner, 2018; Hall and Warner, 2020). Moreover, nest temperatures are substantially cooler for eggs laid earlier (e.g. March – April) vs later (e.g. July – August), resulting in season-specific effects of nest temperature on egg survival and hatchling phenotypes (Pearson and Warner, 2018).

**Figure 1.**
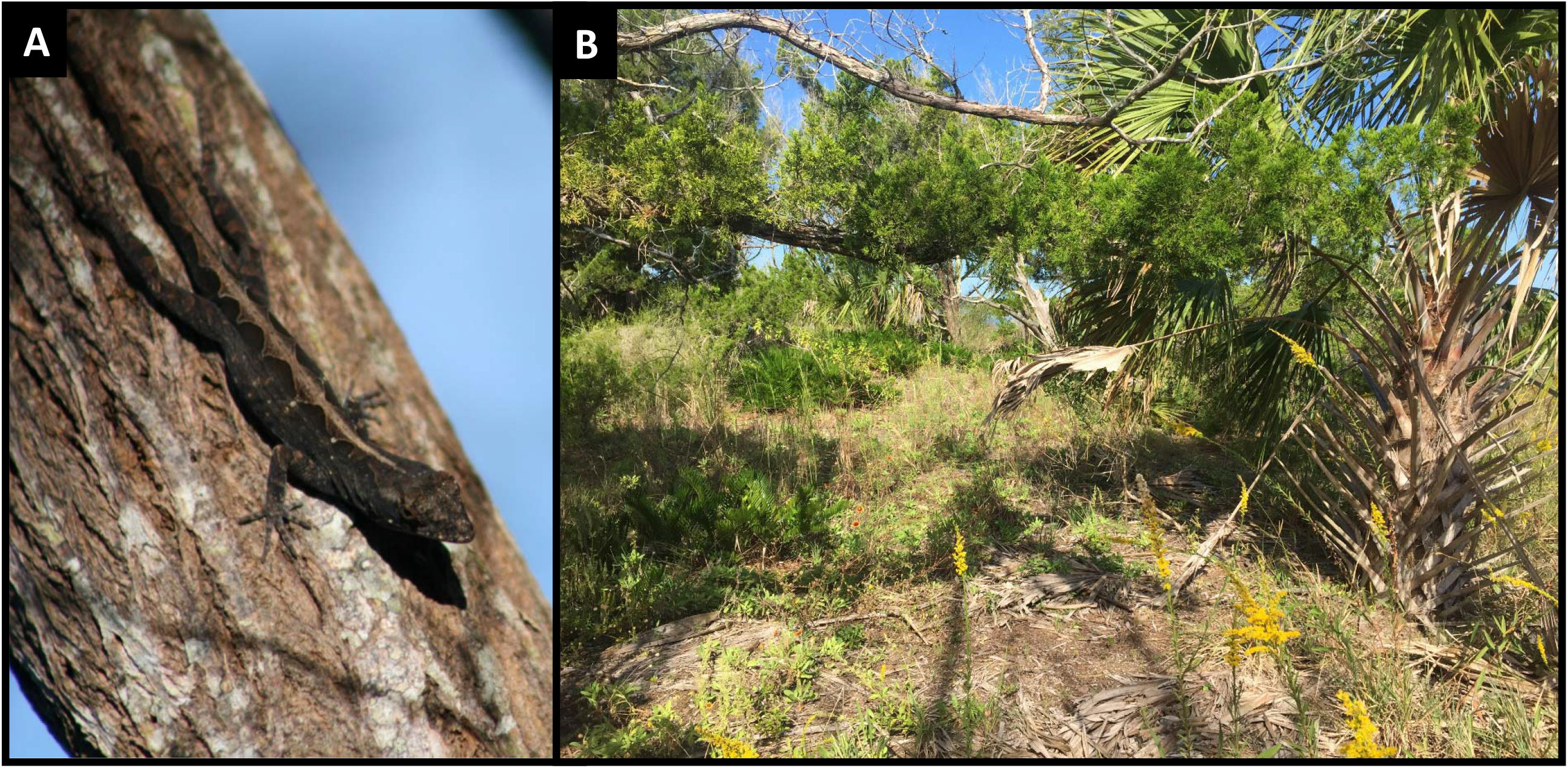
A female brown anole (A) and an example of typical brown anole habitat at our field site in Florida, USA (B).

We captured adult male (49-65 mm SVL; n= 25) and female (42-52 mm SVL; n= 100) *A. sagrei* during 22-23 June 2018 from Palm Coast, FL (29.602199, −81.196211). Lizards were transported to Auburn University to form a captive breeding colony. We housed females individually, in plastic cages (29×26×39 cm) illuminated with Reptisun 5.0 UVB and Tropic Sun 5500K daylight bulbs (Zoo Med Inc., San Luis Obispo, CA, USA) set to a 14 h: 10 h light:dark cycle. Mean cage temperature was 27.6 °C and, due to light sources, oscillated daily from 26 to 30 °C. Cages included reptile cage carpet (Zoo Med Inc.) as a substrate, two bamboo perches, an artificial plant, and a nesting pot (i.e. a plant pot filled with potting soil and peat moss and covered with an artificial leaf). Males were rotated between the same 4 cages of females every 3 days for the first two weeks and then every two weeks thereafter to ensure insemination. However, because we captured females in the middle of the breeding season and they can store sperm for many weeks, all females were likely gravid at time of capture. We fed lizards 4 crickets each, dusted with vitamins and calcium, twice per week and misted cages with water daily.

### Egg collection and treatment allocation

We collected eggs (n= 415) daily from 25 June to 5 August, recorded their mass (to 0.0001 g), date of oviposition, and maternal identity, and randomly distributed them among 8 incubation treatments (see below). Eggs were placed individually in Petri dishes (60 mm x 15 mm) that were half-filled with moist vermiculite (−150 kPa) and wrapped with parafilm to prevent desiccation and control moisture across treatments. To evenly distribute each female’s eggs across treatments, we randomly allocated her first egg to a treatment and then randomly allocated each additional egg to a remaining treatment. No female had more than one egg per treatment.

### Creation of incubation treatments

We used nest temperatures from Pearson and Warner (2018) to create our incubation regimes. Briefly, 22 Thermochron iButtons were placed in various nesting microhabitats at our field site and recorded temperatures every 2.5 hours from 28 March to 15 October 2013. We averaged each 2.5 hour timepoint across iButtons to create two naturally fluctuating regimes: one cooler regime used temperatures from 28 March to 5 May (henceforth “early season”) and one warmer regime used temperatures from 27 June to 4 August (henceforth “late season”). These season-specific incubation temperatures induce variation in egg and hatchling phenotypes (Pearson and Warner, 2018). We refer to these henceforth as the “natural” treatments. Importantly, each natural treatment represents only one potential replication of field temperatures. Indeed, the 22 nests studied exhibit a wide diversity in both mean and thermal variance (Figure S1). Natural nest fluctuations are not symmetrical (i.e. not a sine curve); thus, eggs typically spend more time each day at minimum than maximum temperatures.

Thus, our natural regimes better mimic minimum vs maximum nest temperatures (see Figure S1). Accurately representing both mean and variance of nest temperature in laboratory studies is notoriously challenging (Georges 1989; Georges et al., 2004) and is a common limitation. Regardless, our natural treatments have similar mean temperatures to field nests and include a range of temperatures that represent the majority of those recorded from the field (Figure S1). Moreover, unlike commonly used incubation treatments (described below), our natural treatments allow temperatures to fluctuate widely and stochastically, like real nests.

We used the mean temperatures and variance from these natural treatments to create 6 additional treatments: two constant temperature treatments, two sine fluctuations, and two hourly means fluctuations. These treatments represent commonly used incubation methods in studies of reptile developmental plasticity (While et al., 2018; Booth, 2018). The constant temperature treatments were the raw average of the early and late season natural treatments (Figure 2A). For the sine wave, we created a daily repeating sine fluctuation with an amplitude equal to the mean range of the natural regimes (2.4 °C; Figure 2B). For the repeated hourly means, we took the average temperature at each hour of the day across all iButtons for each season (Figure 2C). Thus, for each season (early vs late) there were 4 methods, resulting in 8 thermal treatments (2×4 factorial design; Figure 2). Treatments were programmed into 8 Memmert IPP55 plus incubators (i.e. 1 incubator per season by treatment incubation regime). Ideally, multiple incubators would be used per season by incubation treatment (i.e. at least 16 incubators; Greenspan et al., 2016); however, this was not possible. There was, however, replication of incubators with respect to season (n = 4 per season) and treatment (n = 2 per treatment).

**Figure 2.**
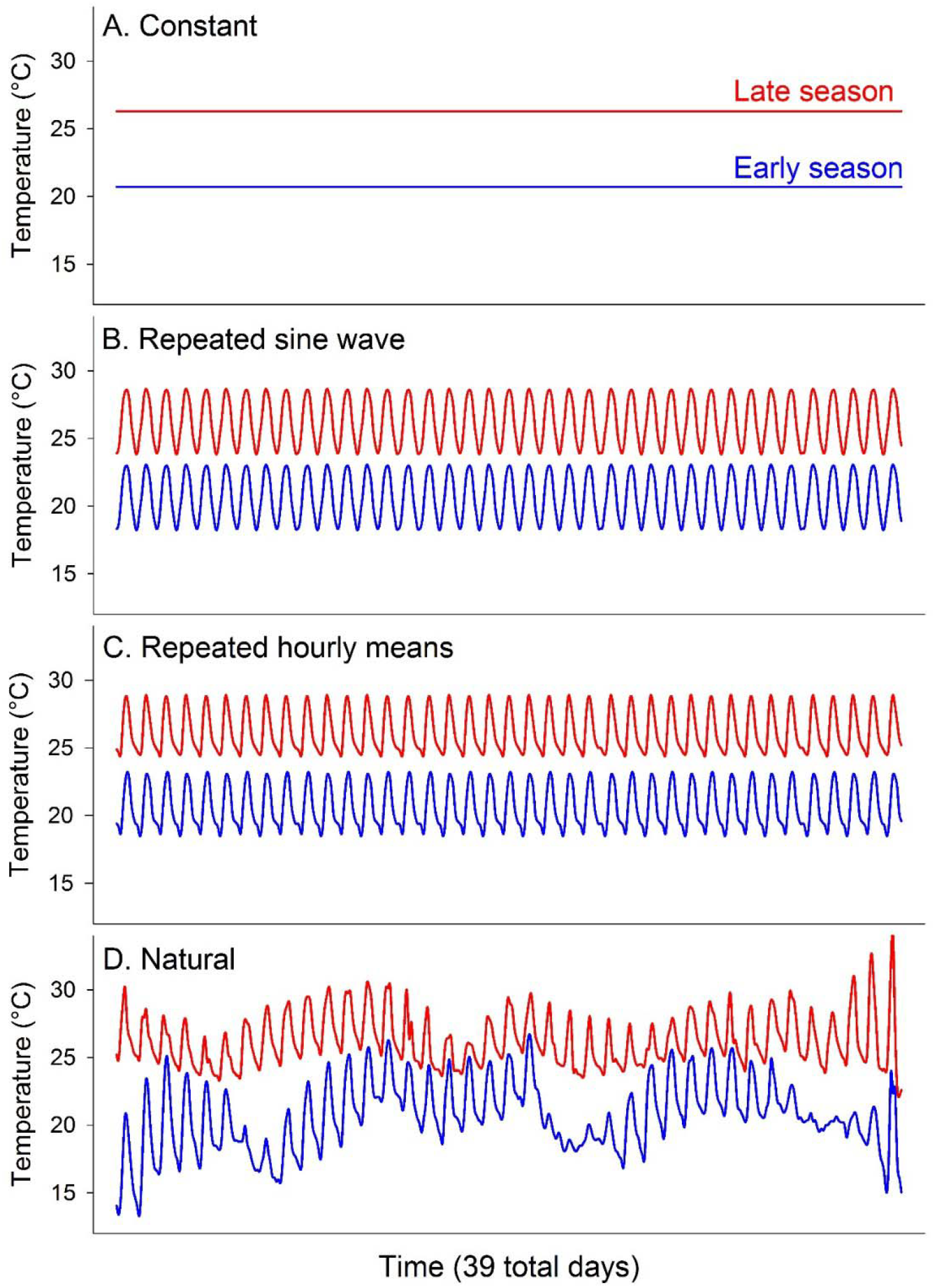
Incubation treatments. Blue and red lines show early- and late-season temperatures, respectively. Each rise and fall of temperature is a daily fluctuation. See Figure 3 for comparison of panels B and C.

The sine and hourly means treatments were programmed to loop daily for the duration of the study. The two natural regimes (early vs late) were each 39 days long, so we programmed incubators to loop this 39-day cycle. Due to temporal variation in egg production, not all eggs in the natural treatments experienced the same temperatures; however, this is a necessary consequence of utilizing a natural thermal regime and is what eggs experience in the wild. All early-season treatments had the same mean temperature (20.7 °C) and all late-season treatments had the same mean temperature (26.3 °C). The sine and hourly mean fluctuations treatments had the same daily temperature range (4.8 °C), which was equal to the mean daily range of temperatures for both the early and late-season natural treatments; however, maximum and minimum daily temperatures and the number of hours spent above and below the mean temperature differed (Figure 3). Thus, within seasons, all treatments had the same means and all fluctuating treatments had the same amount of daily thermal variation but differed according to daily maximum and minimum temperatures. The thermal characteristics used here are important for ectotherm development (Georges et al., 2005), and they allowed critical tests of season by method interactions on offspring traits (i.e. context-dependent effects of methodology).

**Figure 3.**
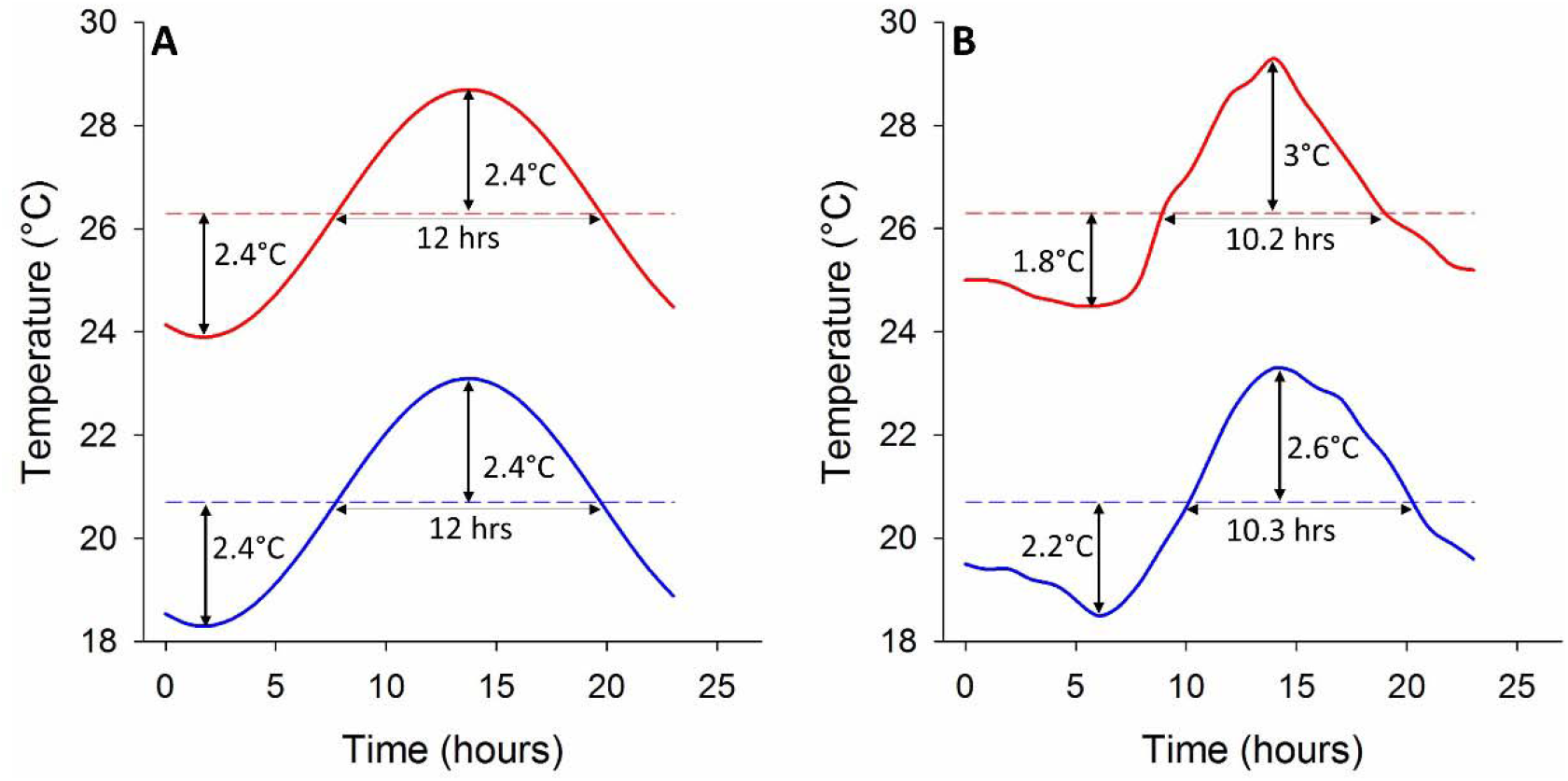
Comparison between sine waves (A) and hourly means (B). Blue and red lines show the early- and late-season temperatures, respectively. Temperature values show the difference between the maximum and minimum temperatures of the fluctuation and the mean temperature (i.e. broken lines). Time values show the total time the fluctuation was warmer than the mean temperature.

### Developmental variables

We measured developmental rate, water uptake, oxygen consumption 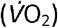, and heart rate (*f*_H_) to assess effects on embryo physiology. We calculated developmental rate by taking the reciprocal of the incubation period (i.e. number of days from oviposition to hatching) and multiplying by 1000 (Viets et al., 1993). For 12 eggs per treatment, we evaluated water uptake by subtracting egg mass at oviposition from egg mass at approximately 65% of development (estimated from Pearson and Warner, 2018). Water uptake during incubation is vital for successful development and has fitness-relevant effects on hatchling phenotypes (Janzen, 1993).

For another 12 eggs per treatment (n = 96 total), we measured 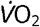 at two temperatures (mean seasonal temperatures: 20.7 and 26.3 °C) at ~ 72% development completed. We used dynamic injection to measure 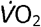 of eggs using a Qubit Q-box RP1LP respirometer (Qubit Biology Inc., Kingston, ON; see Lighton, 2008, pgs 29-31). Each egg was placed in a 10 ml syringe with Luer-Lok tip attached to a 3-way stopcock (Becton, Dickinson and Company, Franklin Lakes, NJ 07417). A 5 μl drop of tap water was placed inside the syringe via a micropipette to prevent desiccation. The plunger was inserted so the volume of the chamber was 6 ml. The stopcock was attached to the respirometer, and the syringe was flushed with CO_2_-free room air for 2 minutes at a rate of 100 ml/min. Two previously drilled holes (between the 5- and 6-ml marks) allowed air to exit the syringe (see Figure 4.3 in Lighton, 2008). After flushing, the syringe plunger was moved down to the 4 ml mark and the stopcock was twisted, sealing the egg in the syringe in a 4 ml volume of air (minus the volume of the egg and the drop of water). The time was noted, and eggs were placed in a constant temperature incubator set to the target temperature (either 20.7 or 26.3 °C) for 3 hours. At the end of 3 hours, the syringe was removed from the incubator, a needle was affixed to the stopcock and a 2 ml sample of air was injected into an injection port in the respirometer. The sample bolus was injected into a stream of dry, CO_2_-free air flowing at 50 ml/min. The time of injection was noted so we could calculate the exact time (in seconds) that each egg was respiring. Control syringes (containing a drop of water but no egg) were treated identically to those described previously and were injected into the respirometer at the beginning and end of the experiment.

All eggs produced from 30 July to 5 August were used to assess embryo *f*_H_ after completing approximately 70% of development (n = 80 eggs; 10 per treatment). For each egg, we measured embryo *f*_H_ at 23 °C using the Buddy^®^ egg monitoring system (Hulbert et al., 2017). Because temperatures fluctuated by 0.5 °C, we used a thermocouple to measure the temperature inside the heart rate monitor for each measure of *f*_H_.

### Post-hatching variables

To assess effects on morphology, we measured the snout-vent length (SVL), body mass, body condition, and tail length of every hatchling (n= 395). Body condition was each hatchling’s residual score from a regression of log mass and log SVL. Thus, higher residuals represent individuals that are relatively heavy for their body length. To assess effects on performance, we measured sprint speed and endurance for approximately 35 hatchlings per treatment (sprint speed – range: n= 31-43 per treatment, total sample size: n = 287; endurance – range: n= 30-47 per treatment, total sample size: n= 298) with a circular racetrack. The racetrack consisted of two circles of aluminum flashing on top of a wooden panel. The outer and inner pieces of flashing were 1.35 and 1.06 meters in circumference, respectively. The racetrack was heated from underneath to keep it within the range of preferred body temperatures for *A. sagrei*. We used a thermocouple to measure the surface of the racetrack during each trial (range= 28.8 – 33.1 °C; mean= 31.2 °C). Lizards were raced within 2 days of hatching: after hatching, each lizard was placed in a cage (see below) and misted with water. On the following day, lizards were raced. No food was provided prior to racing. For racing, hatchlings were placed individually in 50 mL centrifuge tubes wrapped with duct tape to prevent them from seeing out (to reduce stress). Tubes were placed on the racetrack for 45 minutes to allow the lizard’s body temperature to equilibrate with the racetrack. Lizards were quickly placed on the racetrack, one at a time, and chased in a circle using a paintbrush until exhaustion. Exhaustion was the point at which we tapped the lizard on the base of the tail 11 times without it making any movement. All racing trials were conducted by JMH. We recorded the total distance each lizard ran and the total time for each trial. To calculate total distance, we considered the distance of one lap to be the circumference of the center of the raceway (1.21 m). Each hatchling was raced once. The total distance each hatchling ran was our measure of endurance. Sprint speed for each lizard was calculated as the distance of the first lap divided by the time it took to complete that lap.

We measured hatchling growth and survival in the lab over 36 days for 25 hatchlings per treatment. Starting on 6 August, we kept every hatchling until we reached our target sample size for each treatment. Hatchlings were raced and then housed individually in plastic cages (13 x 21 x 17 cm). Each cage contained reptile carpet (Zoo Med Inc.) as a substrate, several plastic leaves, and one wooden perch. Lighting sources, light cycles, and cage temperatures were as described for adults. We provided each lizard 20 fruit flies dusted with vitamins and calcium twice weekly and misted cages with water daily. At the end of 36 days, we measured the SVL of survivors (i.e. Final SVL).

All egg and hatchling phenotypes were selected because 1) they are commonly measured phenotypes in studies of reptile developmental plasticity and 2) they are sometimes correlated with fitness (Booth, 2018; While et al., 2018).

### Statistical analysis

All analyses were performed in R (R Core Team, 2018). Due to variation in egg production among females, the number of females that contributed eggs to each analysis ranged from 63 to 95 (mean 84.8) with some females contributing only 1 egg and others contributing as many as 7 (see Table S1). Therefore, to assess the potential for maternal effects to influence results, for each response variable, we compared two models with a likelihood ratio test: one that included maternal ID as a random effect and one that did not (Table S2). In all models, fixed effects were season, treatment, and their interaction. For some analyses, we included appropriate covariates (see Results). Once the best model was selected, we dropped covariates and interaction terms that were not statistically significant (p > 0.05). Because developmental rates differed between seasons and treatments (see Results) and because eggs often varied in age at time of measurements due to the nature of egg production (i.e. single-egg clutches), we calculated a developmental age to use as a covariate for some analyses (see Results). To calculate developmental age, we divided the age (days since oviposition) at the time an egg was measured by the mean incubation period (in days) for its treatment then multiplied by 100.

For egg and hatchling survival, we conducted generalized linear models with a binomial distribution using the ‘stats’ package in R (i.e. “glm” function) and mixed effects models using the ‘lme4’ package (i.e. “glmer” function; Bates et al., 2007). For developmental and hatchling variables we conducted general linear models using the ‘stats’ package in R (i.e. “lm” function) and linear mixed models using the ‘lme4’ package (i.e. “lmer” function; Bates et al., 2007). For mixed models, maternal ID was a random effect, and we used the ‘lmerTest’ package (Kuznetsova et al., 2017) to assess statistical significance of fixed effects by calculating denominator degrees of freedom via the Satterthwaite approximation. We used the ‘emmeans’ package (Lenth et al., 2018) to make post-hoc comparisons and adjusted p-values via false discovery rate correction. Assumptions of statistical tests were evaluated by visually inspecting model residuals. We log-transformed developmental rate and hatchling endurance to reduce heteroscedasticity and normalize residuals.

## RESULTS

### Developmental variables

Maternal ID improved model fit for developmental rate and 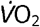 consumption at 26.3 °C (Table S2). In general, the effects of treatment were minor compared to the seasonal effect and only developmental rate was influenced by the interaction (Table 1; Figure 4). Egg survival and water uptake were affected by seasonal temperatures: eggs incubated at early-season temperatures were 2.99 (1.06 – 8.44; 95% confidence limits) times more likely to die and absorbed 93.77 mg (± 10.6 SE) more water than late-season incubated eggs (Table 1,2; Figure 4A,B). Season and treatment interactively influenced developmental rate (Table 1): at early-season temperatures, eggs in the constant treatment had slower developmental rates than all fluctuating treatments and the natural treatment increased developmental rate compared to hourly means (Table 2; Figure 4C). At late-season temperatures, the difference between the constant temperature treatment and natural treatment was not statistically significant, but both resulted in slower developmental rates than the sine and hourly means treatments (Table 2; Figure 4C). There were no statistically significant effects of treatment, season, or their interaction on embryo *f*_H_ or 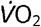 at 20.7 °C (Table 1; Figure 4D,E); however, we did observe a treatment effect on 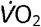 at 26.3 °C (Table 1). Eggs incubated at constant temperatures had greater rates of oxygen consumption than all other treatments; however, the difference between the constant and sine treatments was marginally not significant (p = 0.05; Table 2; Figure 4F). See Table S5 for sample sizes, raw means, and standard deviations of each developmental variable.

**Figure 4.**
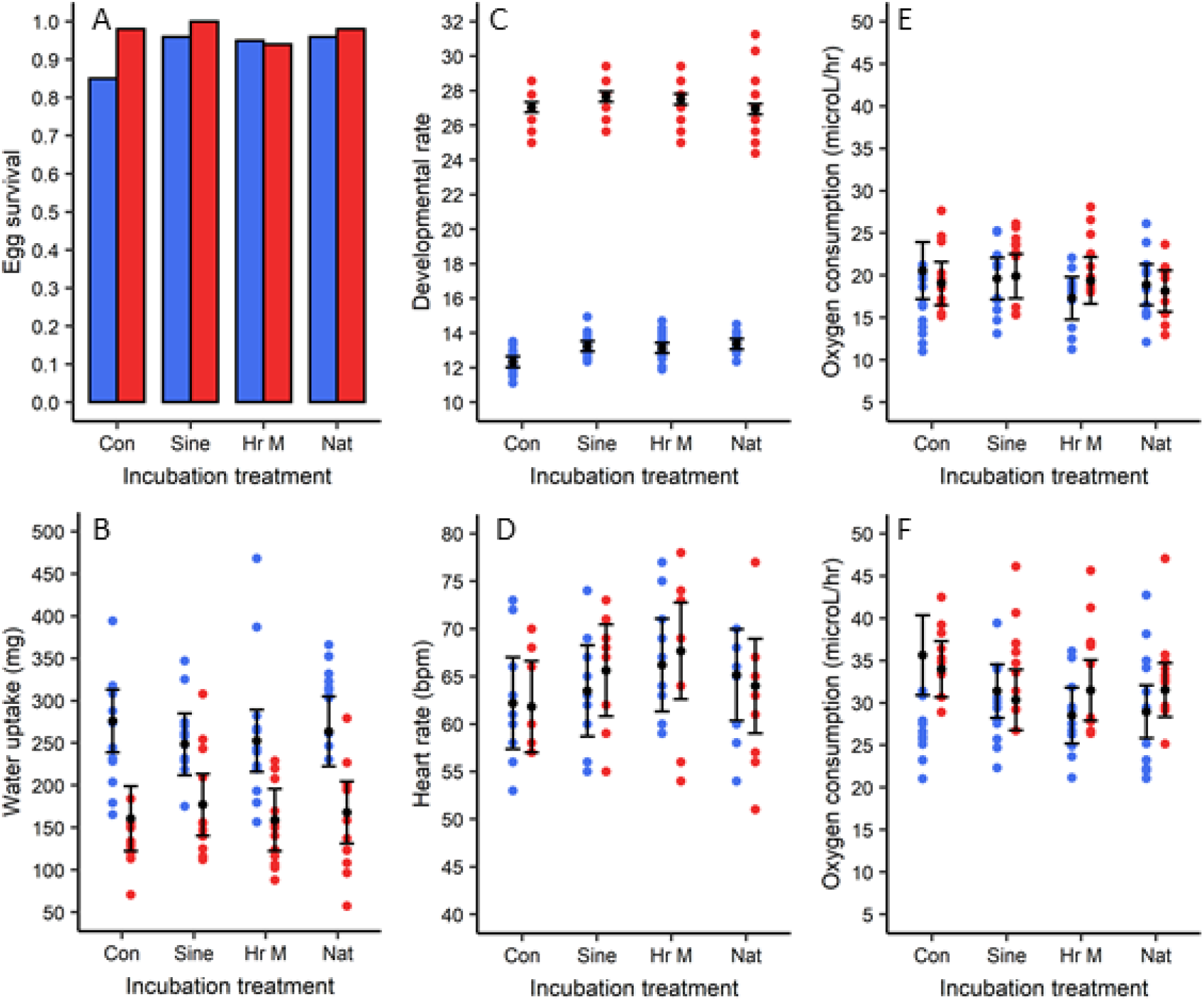
Effects of treatments on developmental variables. Blue and red bars or circles represent early- and late-season temperatures, respectively. Colored circles show raw data, black circles show estimated marginal means (via package ‘emmeans’), and vertical bars show 95% confidence intervals. Con = constant, Sine = sine wave, HrM = hourly means, Nat = natural.

**Table 1.** Results of season, treatment, and their interaction on developmental variables. For covariates, egg mass is mass at oviposition and temperature is the temperature inside the heart rate monitor. See statistical methods for a description of developmental age. See Table S3 for covariate effect sizes. A dash indicates that a covariate or interaction was removed from the model due to a lack of statistical significance. Bold text denotes statistical significance (p < 0.05).

| Response variable | Covariate | Covariate results | Season | Treatment | Season x treatment |
| --- | --- | --- | --- | --- | --- |
| Survival<sup>b</sup> | Egg mass | - | $\chi^2_1=4.89$;<br>**$p = 0.027$** | $\chi^2_3=6.17$;<br>$p = 0.10$ | - |
| Water uptake<sup>b</sup> | Developmental age | **$F_{1,90}=37.01$;**<br>**$p < 0.0001$** | **$F_{1,90}=76.54$;**<br>**$p < 0.0001$** | $F_{3,90}=0.29$;<br>$p = 0.83$ | - |
| Developmental rate<sup>a*</sup> | Egg mass | - | **$F_{1,323}=41721.99$;**<br>**$p < 0.0001$** | **$F_{3,328}=34.04$;**<br>**$p < 0.0001$** | **$F_{3,326}=21.93$;**<br>**$p < 0.0001$** |
| Heart rate<sup>b</sup> | Temperature | **$F_{1,73}=10.44$;**<br>**$p = 0.002$** | $F_{1,73}=0.16$;<br>$p = 0.69$ | $F_{3,73}=2.30$;<br>$p = 0.08$ | - |
| O<sub>2</sub> consumption (20.7 °C)<sup>b</sup> | Developmental age | **$F_{1,87}=25.73$;**<br>**$p < 0.0001$** | $F_{1,87}=0.03$;<br>$p = 0.39$ | $F_{3,87}=1.08$;<br>$p = 0.22$ | - |
| O<sub>2</sub> consumption (26.3 °C)<sup>a</sup> | Developmental age | **$F_{1,81}=43.6$;**<br>**$p < 0.0001$** | $F_{1,64}=1.80$;<br>$p = 0.18$ | **$F_{3,65}=3.60$;**<br>**$p = 0.02$** | - |

Asterisks indicate the response variable was log-transformed; Superscripts a and b denote models that did and did not include maternal ID as a random effect, respectively.

**Table 2.** Pairwise comparisons and log odds ratios for developmental variables. Bold text denotes statistical significance (p < 0.05). P values are adjusted via the false discovery rate method. See Table S4 for estimated marginal means of each variable.

| Response | Season | Contrast/ratio | Estimate | SE | Df | Statistic | P |
| --- | --- | --- | --- | --- | --- | --- | --- |
| Survival (odds ratio) | Early / Late | **Early / Late** | **0.33** | **0.18** | Inf | **-2.07** | **0.038** |
| Water uptake (mg) | Early - Late | **Early - Late** | **93.89** | **10.73** | **90** | **8.75** | **< 0.0001** |
| Developmental Rate* | Early | **Con - HrM** | **-0.064** | **0.007** | **336.4** | **-8.571** | **< 0.0001** |
|  | Early | **Con - Nat** | **-0.081** | **0.007** | **325.0** | **-10.831** | **< 0.0001** |
|  | Early | **Con - Sine** | **-0.071** | **0.007** | **322.56** | **-9.766** | **< 0.0001** |
|  | Early | **HrM - Nat** | **-0.017** | **0.007** | **322.13** | **-2.376** | **0.027** |
|  | Early | HrM - Sine | -0.008 | 0.007 | 324.08 | -1.105 | 0.27 |
|  | Early | Nat - Sine | 0.009 | 0.007 | 320.29 | 1.284 | 0.24 |
|  | Late | **Con - HrM** | **-0.018** | **0.007** | **331.01** | **-2.404** | **0.025** |
|  | Late | Con - Nat | 0.004 | 0.007 | 336.31 | 0.577 | 0.56 |
|  | Late | **Con - Sine** | **-0.024** | **0.007** | **320.44** | **-3.306** | **0.003** |
|  | Late | **HrM - Nat** | **0.022** | **0.007** | **330.18** | **2.97** | **0.006** |
|  | Late | HrM - Sine | -0.006 | 0.007 | 330.01 | -0.82 | 0.50 |
|  | Late | **Nat - Sine** | **-0.028** | **0.007** | **330.29** | **-3.844** | **0.001** |
| O<sub>2</sub> consumption (26.3 °C) | Combined | **Con - HrM** | **4.13** | **1.40** | **76.25** | **2.94** | **0.01** |
|  | Combined | **Con - Nat** | **3.95** | **1.35** | **74.19** | **2.92** | **0.01** |
|  | Combined | Con - Sine | 3.18 | 1.38 | 74.39 | 2.31 | 0.05 |
|  | Combined | HrM - Nat | -0.18 | 1.24 | 77.04 | -0.15 | 0.88 |
|  | Combined | HrM - Sine | -0.94 | 1.22 | 71.80 | -0.78 | 0.60 |
|  | Combined | Nat - Sine | -0.76 | 1.13 | 49.03 | -0.68 | 0.60 |

Con = constant, HrM = hourly means, Nat= natural; SE = standard error; Df = degrees of freedom; Statistic = t statistic or z ratio for contrasts and log odds ratios, respectively; P = p value; Inf = infinity degrees of freedom; Asterisks denote a variable was log transformed

### Post-hatching variables

Maternal ID improved model fit for all measures of hatchling morphology but not for performance, growth, or survival (Table S2). In general, the effects of treatment were minor compared to seasonal effects and only hatchling endurance was influenced by the interaction (Table 3; Figures 5, 6. Late-season temperatures produced shorter (Figure 5A) but heavier offspring (Figure 5B, C), with longer tails (Figure 5D) compared with early-season temperatures (Table 4). Initial analysis revealed that SVL was significantly affected by treatment (Table 3); however, this effect was not statistically significant after p-values were corrected for post-hoc comparisons (Table 4). Body condition was the only morphological variable that was clearly affected by treatment (Table 3). Hatchlings from the hourly means treatment had greater body condition than those from the sine wave and constant temperature treatments, but no other differences were statistically significant (Table 4; Figure 5C).

**Figure 5.**
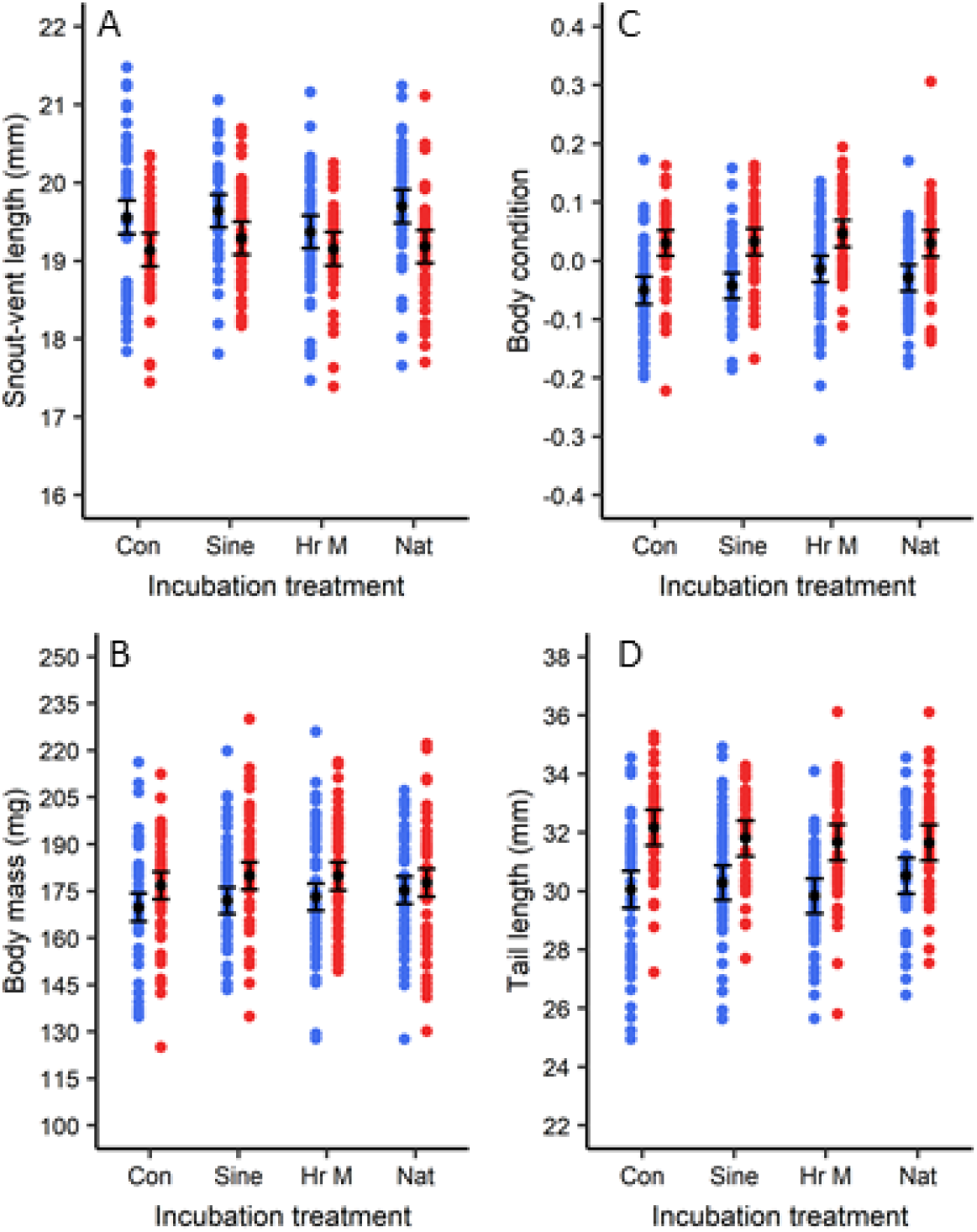
Effects of treatments on hatchling morphology. Blue and red circles represent early-and late-season temperatures, respectively. Colored circles show raw data, black circles show estimated marginal means (via package ‘emmeans’), and bars show 95% confidence intervals. Con = constant, sine = sine wave, HrM = hourly means, Nat = natural.

**Table 3.**
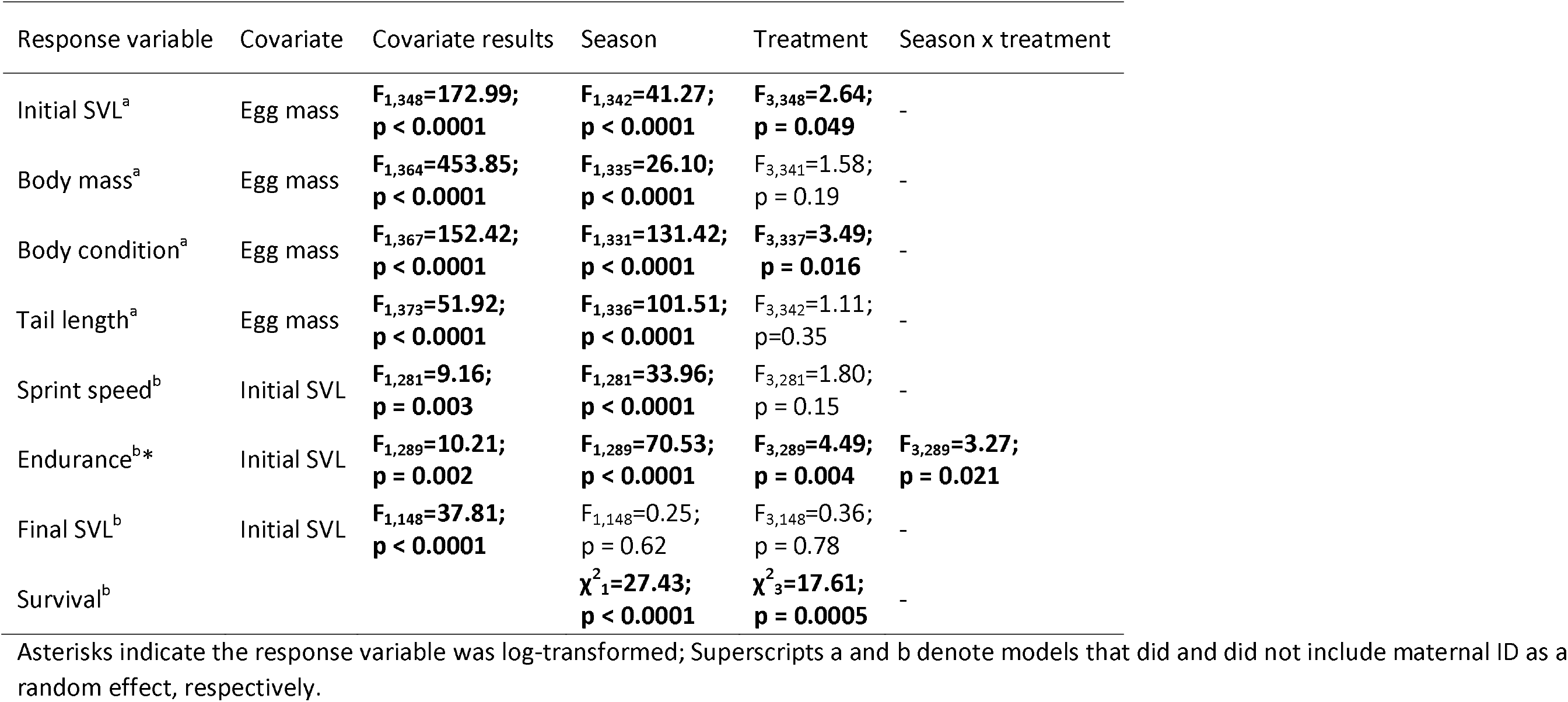
Results of season, treatment, and their interaction on post-hatching variables. For covariates, egg mass is mass at oviposition and initial SVL is SVL at time of hatching. No covariate was considered for analysis of survival. See Table S6 for covariate effect sizes. A dash indicates that the interaction was removed from the model due to a lack of statistical significance. Bold text denotes statistical significance (p < 0.05).

Sprint speed was greater for hatchlings from late-season temperatures (Table 4; Figure 6A), but neither treatment nor the interaction were statistically significant (Table 3). We observed an interaction for hatchling endurance: at early-season temperatures, hatchlings from the natural treatment had greater endurance than those from the constant temperature and sine treatments and those from the hourly means treatment had greater endurance than those from the constant treatment; however, there were no significant treatment effects for late-season incubated eggs (Table 4; Figure 6B). There were no statistically significant effects of season, treatment, or their interaction on hatchling final SVL, indicating that growth did not differ among treatments (Table 3; Figure 6C).

**Figure 6.**
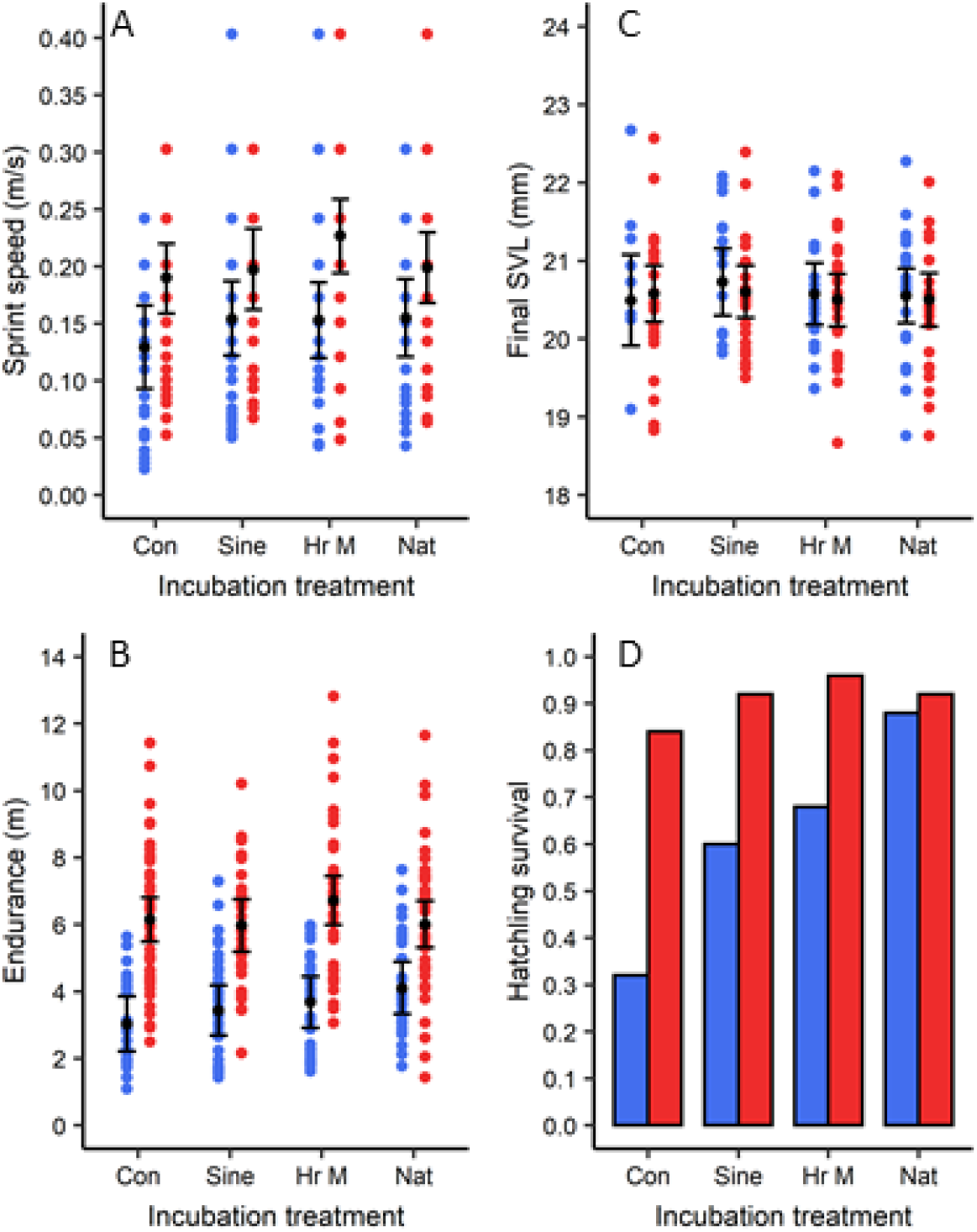
Effects of treatments on post-hatching variables. Blue and red bars or circles represent early- and late-season temperatures, respectively. Colored circles show raw data, black circles show estimated marginal means (via package ‘emmeans’), and horizontal bars show 95% confidence intervals. Con = constant, Sine = sine wave, HrM = hourly means, Nat = natural.

**Table 4.**
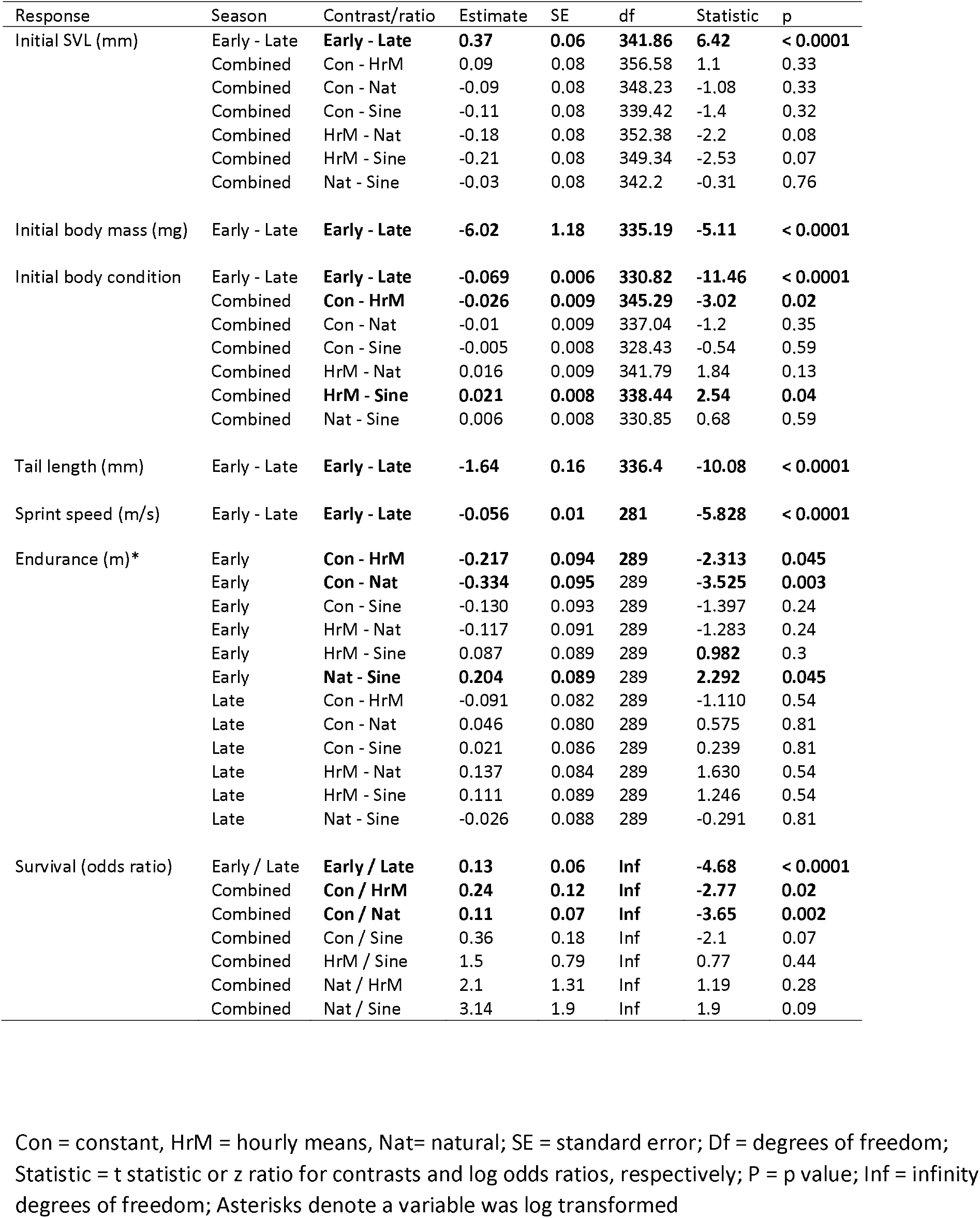
Pairwise comparisons and log odds ratios for post-hatching variables. Bold text denotes statistical significance (p < 0.05). P values are adjusted via the false discovery rate method. See Table S7 for estimated marginal means of each variable.

We observed season and treatment effects on hatchling survival, but the interaction was not statistically significant (p = 0.42; Table 3). Hatchlings from early-season incubation temperatures were 7.54 (3.24 – 17.5; 95% confidence limits) times as likely to die as those from late-season temperatures (Table 4); however, this was primarily driven by high mortality in the early-season constant, sine, and hourly means treatments (Figure 6D). Hatchlings from the natural and hourly means treatments were 8.70 (1.9 – 39.93; 95% confidence limits) and 4.15 (1.11 – 15.51; 95% confidence limits) times more likely to survive than those from the constant temperature treatment, respectively, but no other differences were statistically clear (Table 4; Figure 6D). See Table S8 for sample sizes, raw means, and standard deviations of each post-hatching variable.

## DISCUSSION

Several studies compare effects of constant vs repeated temperature fluctuations on development (e.g. birds [Olson et al., 2006], lizards [Du and Shine, 2010], frogs [Niehaus et al., 2006; Arrighi et al., 2013], fish [Eme et al., 2018]; insects [Liu et al., 1995; Folguera et al., 2011; Xing et al., 2019]). A few studies measure or model the effects of natural thermal regimes on biological processes (e.g. Niehaus et al., 2012; Bannerman and Roitberg, 2014) or test differences in constant temperature, repeated daily fluctuations, and random changes in diurnal fluctuations (e.g. Schaefer and Ryan, 2006; Manenti et al., 2014). To our knowledge, however, no study of developmental plasticity has compared constant temperature and repeated fluctuations to natural nest fluctuations under different contexts (e.g. season) in the laboratory. Some have compared eggs incubated in the lab and in the field (e.g. St Juliana and Janzen, 2007; Paitz et al., 2010); however, these studies cannot pinpoint thermal effects because there are many other variables in natural nests that influence development (Warner et al., 2018).

### Developmental variables

Seasonal effects on development were typical: warmer temperatures increase rates of survival and development and reduce water uptake (Gutzke and Packard, 1987; Pearson and Warner, 2018). The absence of a season effect on 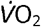 and *f*_H_ indicates there was no thermal acclimation of embryo physiology which is typical for shallow-nesting species (like anoles) but not for species that nest deeper in the soil (Du et al., 2010). The effects of incubation treatment were minimal for most egg phenotypes, but some fitness-relevant variables were affected (e.g. developmental rate). Moreover, treatment effects were small compared to those of seasonal temperature. In studies of thermal developmental plasticity, the largest effects are often on developmental rates (Stillwell and Fox, 2005; Noble et al., 2018), and our data indicate this is also true for incubation methods. Developmental rates can have important effects on fitness, given that early hatching is often associated with increased hatchling survival (Pearson and Warner, 2018) and faster development reduces embryo exposure to adverse conditions (Doody, 2011). At cooler temperatures, the natural treatment increased developmental rates relative to other treatments; however, at warmer temperatures, the natural treatment somewhat slowed development relative to other treatments. This likely results from the curvilinear relationship between developmental rates and temperature (see Fig. 3 in Bowden et al., 2014): at early-season temperatures, relatively small reductions in developmental rate due to cold temperatures are offset by relatively greater increases at warmer temperatures and vice versa for the late-season treatment. This is particularly important for species with temperature-dependent sex determination (TSD), where the proportion of development spent at a given temperature determines sex (Georges 1994). Thus, slight differences in incubation methods could influence sex ratios (Neuwald and Valenzuela, 2011; Bowden et al., 2014) and fitness (Warner and Shine, 2011). Studies that compare the different incubation methods used here to examine sex ratios in species with TSD would provide further insight into the ecological-relevance of the effects of lab incubation on this important demographic trait.

Intriguingly, constant temperature treatments resulted in greater metabolic rates than other treatments. There are biological and methodological explanations for this result. Embryo metabolic acclimation to temperature is widespread across reptiles and likely serves to compensate for environmental conditions that reduce development rates (e.g. cool temperatures; Du et al., 2010). Constant temperatures result in relatively slow development compared to fluctuating conditions. Thus, embryos may compensate by upregulating metabolism when temperatures are thermostable. We think this biological explanation is unlikely because we did not observe metabolic acclimation with respect to temperature (i.e. early vs late treatments). An alternate explanation is that embryos in the constant treatment were younger and smaller at the time 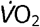 was measured due to treatment specific variation in developmental rates. Because 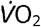 scales with embryo size and, thus, age, using developmental age as a covariate may have caused the model to overestimate mean values of 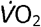 for the constant treatment. Note that the 95% CI of the estimated marginal mean 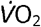 for the early-season constant treatment excludes nearly all the raw data in Figure 4F. Although several studies have examined metabolic acclimation of embryos to temperature (see Du and Shine, 2015), to our knowledge, no study has explicitly considered the effects of fluctuating vs constant temperatures on acclimation.

### Post-hatching variables

Seasonal effects on post-hatching variables were typical: warmer temperatures have small effects on SVL and hatchling growth but increase body mass, body condition, tail length, performance, and survival in the laboratory (Pearson and Warner, 2018). Treatment effects on hatchling phenotypes were relatively small and most were not statistically clear. However, there was a relatively large effect on hatchling survival – a critical component of fitness. Although the interaction between season and treatment was not statistically significant, the natural regime vastly improved hatchling survival compared to other early-season treatments (i.e. the effect was large; Figure 6D). This may be, in part, related to performance since, at colder temperatures, hatchlings from the natural treatment had greater endurance than other groups (Table 4). Additionally, sprint speed and growth rates were lowest for hatchlings from the early-season constant temperature treatment, though these effects were not statistically significant. In general, hatchling performance and survival are lower at extreme incubation temperatures compared to intermediate temperatures which are within the optimal thermal range (i.e. OTR; Andrews and Schwarzkopf, 2012; Noble et al., 2018). Although eggs in the early-season, natural treatment briefly experienced temperatures as low as 13 °C, there were many days that temperatures peaked at 24-26 °C (Figure 2D), which is well-within the OTR for this species (i.e, 22 – 29 °C; Hall and Warner, 2020). Because developmental rates increase with temperature, in the early-season natural treatment, eggs spent a greater proportion of development at temperatures within the OTR than in other treatments, and this may have enhanced hatchling performance and survival. Evidence for this speculation is that, for early-season temperatures, the natural treatment resulted in a faster developmental rate than all other treatments (Table 2). Therefore, at colder temperatures, using a naturally fluctuating regime may optimize development compared to repeated fluctuations or constant temperatures, whereas at warmer temperatures, which are within the OTR, it may have little effect.

There are two important conclusions to draw from the hatchling survival results. First, these effects must have resulted from treatment specific patterns of embryo development since the treatments were applied to eggs. Thus, conditions appropriate for successful embryo development may not necessarily be best for hatchlings (e.g. compare early-season constant egg vs hatchling survival). Consequently, researchers should assess the effects of egg environments on hatchling performance and survival. Although assessing the influence of developmental environments on later life stages will be a critical advancement (Refsnider et al., 2019), the importance of this could depend on whether research goals are focused on proximate mechanisms of thermal effects vs the fitness consequences. Second, although incubation treatments had minimal effects on egg phenotypes and hatchling morphology, they had relatively large effects on performance and survival (i.e. fitness), indicating that many commonly measured phenotypes may not be good fitness proxies. Although numerous studies demonstrate that incubation temperature influences locomotor performance in reptiles, very little work has been done to unearth the mechanisms driving these patterns (Booth, 2018). Most likely, thermal effects on performance are driven by morphological and physiological variation at cellular and subcellular levels (e.g. number and type of muscle fibers; mitochondrial density; oxygen transport; reviewed by Booth, 2017, 2018); therefore, we should not necessarily expect gross measures of hatchling morphology to coincide with performance and fitness. Our study adds a new layer of complexity by demonstrating that the mean and variance of incubation temperature can potentially interact to determine locomotor performance and survival, independent of gross morphology (e.g. body size).

### Implications for experimental design and study comparisons

Most studies that incorporate fluctuating temperatures use repeated sine waves (e.g. Folguera et al., 2011; Xing et al., 2019), few use an hourly means fluctuation (e.g. Davis et al., 2006; Tiatragul et al., 2017; Hall and Warner, 2018) and almost none use a natural regime (Pearson and Warner, 2018; Tiatragul et al., 2020). The largest effects of incubation method were, generally, between constant and fluctuating treatments. For example, the effects of sine waves or hourly means were similar and neither differed substantially from the natural treatment for most phenotypes. Thus, researchers should use fluctuating rather than constant temperatures; however, the type of fluctuation may be of little importance if temperatures are within the OTR. Sine waves are particularly advantageous because they are easy to standardize and replicate. Conversely, recreating natural regimes is costly with respect to equipment and time. For example, to create our natural treatments, we made programs with 39 different fluctuations, and, due to limited memory in incubators, these were assimilated into 6 individual programs that were each manually uploaded approximately once per week throughout the study. This extra effort is only worthwhile under certain contexts (i.e. colder temperatures). If we had conducted a study using only early and late season constant temperatures (or repeated sine waves) we would have concluded that early-season temperatures are less conducive to proper development due to relatively large treatment effects on hatchling survival; however, using natural fluctuations demonstrates that early and late temperatures each result in high hatchling survival (Figure 6D).

Finally, great effort is given to describing general trends in ecological and evolutionary phenomena (e.g. meta-analyses and literature reviews on reptile developmental plasticity: Noble et al., 2018; Warner et al., 2018; Mitchell et al., 2018; While et al., 2018; Refsnider et al., 2019). Such broad syntheses of published data depend on methodological consistency across studies and repeatability of results. We found mixed support for the assumption that data are comparable across incubation methods. For example, increasing thermal complexity (from constant to natural treatments) had little influence on egg survival, but it vastly improved survival of hatchlings at colder temperatures. Due to these context-dependent effects, researchers should exercise caution when making comparisons across studies that utilize both different methods and temperatures and seriously consider the thermal physiology of their organism along with methodology.

### Biological relevance vs statistical significance

Due to the misuse and misinterpretation of null hypothesis significance testing in ecology and evolution, researchers must always evaluate biological relevance and statistical significance (Martinez-Abrain, 2008). Our results afford opportunity to discuss this issue with respect to incubation methods. Slight differences in treatment occasionally resulted in statistically significant, but potentially biologically trivial differences among groups. For example, hatchlings from early-season incubation temperatures were longer in SVL than late-season incubated hatchlings; however, this extra length only equates to a 1.9% difference. Additionally, it did not afford greater locomotor performance to early-season incubated hatchlings, even though longer lizards, in general, run faster and have more endurance (Table S6). Moreover, past studies using similar incubation temperatures show no effect of temperature on SVL (Pearson and Warner, 2018) or the opposite effect of that observed in our study (Pearson and Warner, 2016), indicating our result may be spurious.

Conversely, treatment effects on hatchling survival were much greater at early vs late season temperatures, even though the interaction term was not statistically significant (Figure 6D). Given the large effect size, ignoring this potential interaction may lead to false conclusions. Indeed, power for detecting interactions is low for binomial data even when sample sizes are relatively large (i.e. greater type II error rate; Marshall, 2007). Normally, we would not conduct post-hoc tests in the absence of a statistically significant interaction; however, we do so here to emphasize biological relevance vs statistical significance. Post-hoc inspection indicates that the natural and constant treatments differ for early but not late season temperatures (Figure S2; Figure S3) and that early season temperatures resulted in lower survival than late season temperatures for constant, sine, and hourly means treatments but not the natural treatment (Figure S4). Therefore, despite a lack of statistical significance, the biological relevance seems clear: natural fluctuations greatly enhance offspring survival at cooler temperatures, particularly in comparison to constant temperatures; however, at warmer temperatures, thermal fluctuations are less important (Figure 6D). Ultimately, these data indicate that considering biological relevance and statistical significance is vital when comparing results across studies that use different incubation methods, even if mean temperatures are comparable.

### Limitations and caveats

One important limitation is that our study species constructs shallow nests that are prone to extreme thermal fluctuations (Hall and Warner, 2020). Many species, however, construct nests in thermostable environments and constant incubation temperatures are likely optimal for development (e.g. Andrews, 2018). Thus, our results are not generalizable across some species. Another caveat is that we used a single natural regime per seasonal temperature to represent “natural” conditions, when there are large differences in mean and variance of temperature across natural nests (Figure S1). Moreover, our natural regimes were created from taking mean temperatures across nests, which reduces the thermal variation present in nature. Alternate natural regimes that replicate more extreme conditions may generate different results. A future study could potentially improve upon our methods by replicating the coldest and warmest nests in addition to a nest of intermediate temperature (rather than use mean temperatures); however, experimenters are always limited with respect to resources (e.g. incubators) and sample sizes. We emphasize that this experiment required approximately 400 eggs, which limits the number of viable study species and illustrates the difficulty of utilizing an even more complex design (e.g. multiple natural treatments per season).

### Conclusions

The degree to which greater levels of treatment complexity influences results is dependent on ecological factors important to the study system, and researchers should invest time in evaluating methods in a context-dependent way. Our results are important for all researchers attempting to achieve greater levels of ecological relevance in studies of thermal ecology. Our design can determine how much thermal complexity is required to effectively reproduce real thermal environments under different conditions; thus, minimizing logistical costs of future studies and improving the reliability of results. This work has several important implications for controlled laboratory or field studies of ecology and evolution. First, quantifying the effects of various methods should be done under different ecological contexts so researchers can assess the effects relative to ecological factors important to the study system. Second, depending on the context, using potentially costly, but ecologically-meaningful treatments may illuminate mechanisms that drive biological phenomena or may be unnecessary. Third, the context-dependent nature of methodology should be considered when synthesizing published studies since even small differences in methods can result in biologically meaningful or, sometimes, statistically spurious differences. Finally, ecological or methodological effects may not be detectable in the stage at which they are applied (e.g. egg), but rather manifest later (e.g. hatchlings). Thus, researchers should measure a broad range of phenotypes at multiple life stages when aiming to assess the effects of developmental environments on fitness.

## Acknowledgements

We thank K. Wilson, C. Reali, A. Dees and A. Stephens for help with animal care, A. Appel for methodological advice, and TS Mitchell for helpful comments on an early draft. Research was approved by Auburn University IACUC # 2018-3233 and funded by National Science Foundation (DEB-1564563). J. Hall acknowledges financial support from the Alabama Graduate Research Scholars Program (GRSP) funded through the Alabama Commission for Higher Education and administered by the Alabama EPSCoR. This is publication # 905 of the Auburn University Museum of Natural History.

## Data Accessibility

Data will be available through Auburn University’s digital repository (AUrora).

